# Real Time PCR for the Evaluation of Treatment Response in Clinical Trials of Adult Chronic Chagas Disease: Usefulness of Serial Blood Sampling and qPCR Replicates

**DOI:** 10.1101/343244

**Authors:** Rudy Parrado, Juan Carlos Ramirez, Anabelle de la Barra, Cristina Alonso-Vega, Natalia Juiz, Lourdes Ortiz, Daniel Illanes, Faustino Torrico, Joaquim Gascon, Fabiana Alves, Laurence Flevaud, Lineth Garcia, Alejandro G. Schijman, Isabela Ribeiro

## Abstract

This work evaluated a serial blood sampling procedure to enhance the sensitivity of duplex real time PCR (qPCR) for baseline detection and quantification of parasitic loads and post-treatment identification of failure in the context of clinical trials for treatment of chronic Chagas disease, namely DNDi-CH-E1224-001 (NCT01489228) and MSF-DNDi PCR sampling optimization study (NCT01678599). Patients from Cochabamba (N= 294), Tarija (N = 257), and Aiquile (N= 220) were enrolled. Three serial blood samples were collected at each time-point, and qPCR triplicates were tested per sample. The first two samples were collected during the same day and the third one seven days later.

A patient was considered PCR positive if at least one qPCR replicate was detectable. Cumulative results of multiple samples and qPCR replicates enhanced the proportion of pre-treatment sample positivity from 54.8 to 76.2%, 59.5 to 77.8%, and 73.5 to 90.2% in Cochabamba, Tarija, and Aiquile cohorts, respectively and increased cumulative detection of treatment failure from 72.9 to 91.7%, 77.8 to 88.9%, and 42.9 to 69.1% for E1224 low, short, and high dosage regimes, respectively; and from 4.6 to 15.9% and 9.5 to 32.1% for the benznidazole (BZN) arm in the DNDi-CH-E1224-001 and MSF-DNDi studies, respectively. The monitoring of patients treated with placebo in the DNDi-CH-E1224-001 trial revealed fluctuations in parasitic loads and occasional non-detectable results. This serial sampling strategy enhanced PCR sensitivity to detecting treatment failure during follow-up and has the potential for improving recruitment capacity in Chagas disease trials which require an initial positive qPCR result for patient admission.

## Introduction

Following years of little progress in research and development of new compounds for treatment of Chagas disease (CD), new chemical classes have demonstrated encouraging activity against its causative agent, *Trypanosoma cruzi* [1,2]. The efficacy of anti-*T. cruzi* compounds has habitually been measured by means of parasite detection or antibody titers. However, in chronically infected patients, traditional parasitological methods lack sensitivity and *T. cruzi*-specific antibody titers don’t usually decrease until many years after treatment [2]. In this context, molecular methods, such as conventional and real-time PCR (qPCR) assays, have opened promising opportunities for monitoring bloodstream parasitic levels to detect therapeutic failure or response [3–6]. Following this approach, multicenter PCR studies have allowed harmonization and validation of standard operating procedures (SOPs) for PCR-based detection and quantification of *T. cruzi* DNA in blood samples [7,8] coupled with external control quality assurance [9]. However, the best performing qPCR methods reached between 60-70% of positivity in untreated chronic Chagas disease patients when a single baseline blood sample was tested [7,8,10], a figure which has been verified in different clinical trials [11-13].

In clinical trials in which eligibility criteria for patient enrollment include PCR positivity, such low values of sensitivity require that a larger proportion of seropositive subjects must be screened before being admitted. To overcome this limitation, a PCR sampling optimization study (NCT01678599) was developed by Drugs for Neglected Diseases *initiative* (DND*i*) and Médecins Sans Frontières (MSF) with the aim of evaluating sampling conditions for qPCR monitoring of benznidazole (BZN) treatment, as well as DNDi-CH-E1224-001, a DND*i-*sponsored randomized clinical trial (NCT01489228) to evaluate safety and efficacy of three oral regimens of E1224 in comparison with BZN and placebo, which planned to collect three serial peripheral blood samples from each patient at each follow-up time point and to perform qPCR in triplicate from each blood sample DNA extract.

This report presents the data obtained in these studies, showing an improvement in qPCR clinical sensitivity for both enrollment and detection of treatment failure in adult patients with chronic Chagas disease.

## Methods

### Ethics statement

The clinical trials, including the sampling requirements, were approved by the Ethical Review Boards of Universidad Mayor de San Simón, Fundación CEADES, Hospital Clínic and Médecins Sans Frontières, following the principles expressed in the Declaration of Helsinki. Written informed consent forms were signed by the study volunteers (no minor subjects were included in these trials). All samples were anonymized before being processed.

### Subjects and samples

Subjects and samples were recruited from two different clinical studies:

i) The DNDi-CH-E1224-001 clinical trial (NCT01489228), designed and sponsored by DND*i*, a proof-of-concept double-blinded randomized trial aiming to evaluate the safety and efficacy of three (high, low, and short) oral regimens of E1224, compared to BZN (5 mg/kg/day) and placebo, during 60 days of treatment of adult patients with chronic indeterminate Chagas disease [14].

A total of 560 patients aged 18-50 years and serologically confirmed as having Chagas disease were screened in two study sites of The Platform for a Comprehensive Care of Patients with Chagas disease in Bolivia (one site in the city of Cochabamba (N= 294) and the other in the city of Tarija (N= 257)). Of those screened, 551 patients had PCR results available for analyses, as a total of 9 patients withdrew consent for participation and no PCR sample was obtained (an additional patient withdrew consent only during the baseline visit, thus after a screening sample was obtained). Samples consisted of of peripheral blood mixed with an equal volume of guanidine hydrochloride 6M-EDTA 0.2M pH 8.0 (GEB) buffer. A maximum of three 10 mL GEB samples were collected at baseline: sample 1 (S1) and sample 2 (S2) were collected during the same day and sample 3 (S3) seven days later, but only if DNA extracts from S1 and S2 gave non-detectable results. The qPCR was assayed in duplicate from both S1 and S2 DNA extracts. In cases where both replicates gave non-detectable results, a third replicate was analyzed. When all qPCR replicates from both S1 and S2 gave non-detectable results, S3 was collected and assayed in triplicate. During follow-up, the three GEB samples were collected at each time-point visit (end of treatment [EOT], and 2, 4, and 10 months post-treatment) and qPCR was assayed in triplicate from each S1, S2, and S3 DNA sample.

ii) The MSF-DNDi PCR sampling optimization study (NCT01678599) launched by DND*i* and Médecins Sans Frontières (MSF) aimed to evaluate sampling strategies for qPCR treatment monitoring in adult patients with chronic Chagas disease (with indeterminate or early target organ involvement) treated with BZN (5 mg/kg/day) for 60 days. [15, 16] This study was carried out in 17 communities in the rural locality of Aiquile and did not include a placebo or other comparison treatment group. A total of 220 patients aged 18-60 years with serologically confirmed Chagas disease were recruited. Of these, 205 patients (93.2%) had at least 1 positive PCR at baseline or end of treatment. 201 patients (91.36%) had a positive PCR at baseline. All houses of patients entering the study were subjected to entomological surveillance.

From each seropositive patient, three GEB samples were collected at baseline and at each follow-up visit (EOT, 4, and 10 months post-treatment). S1 and S2 were collected during the same day and S3 seven days later. S1 and S3 consisted of 10 mL of blood, whereas for S2 5 mL was collected; all samples were mixed with an equal volume of guanidine-EDTA buffer. The qPCR was assayed in triplicate from each S1, S2, and S3 DNA sample.

Only patients with at least one positive result out of a maximum of nine qPCR replicates were enrolled in these trials. In both studies, therapeutic failure was defined as the persistence of parasite DNA, detected in at least one qPCR replicate, at any time-point during post-treatment follow-up.

### DNA extraction

The High Pure PCR Template Preparation kit (Roche Diagnostics Corp., Indianapolis, IN) was used to process 300 µL of each GEB sample, which was then eluted in 100 µL elution buffer as previously described [10].

### Quantitative real-time PCR procedure

A duplex qPCR targeted to *T. cruzi* satellite DNA (SatDNA) and an internal amplification control (IAC) were used as previously described [10]. The qPCR reactions were carried out with the use of FastStart Universal Probe Master Mix (Roche Diagnostics GmbHCorp., Mannheim, Germany) with 5 μL eluted DNA in a final volume of 20 μL. Cycling conditions were a first step of 10 minutes at 95 °C, followed by 40 cycles at 95 °C for 15 seconds, and a final step of 1 minute at 58 °C. The amplifications were carried out in a Rotor-Gene Q (Corbett LifeScience, Cambridgeshire, United Kingdom) real-time PCR device.

For quantification purposes, standard curves were plotted with 1/10 serial dilutions of total DNA obtained from a GEB seronegative sample spiked with 10^5^ par. eq./mL LL014-1-R1 Cl1 *T. cruzi* stock (TcV) cultured epimastigotes. A negative control and two positive controls containing 10 and 1 fg/µL *T. cruzi* CL-Brener DNA were included in every run, as recommended [17].

### Genotyping of *T. cruzi* discrete typing units

Baseline samples from both clinical studies with SatDNA qPCR Ct (threshold cycle) values below 33 (N= 180) were genotyped using PCR-based strategies targeted to nuclear genomic markers, namely: (1) spliced leader intergenic region (SL-IR) based PCR was used to distinguish TcI (150 bp), and TcII, TcV, and TcVI (157 bp) from TcIII and TcIV (200 bp); (2) heminested SL-IR-I PCR was used to confirm TcI (350 bp) and heminested SL-IR-II PCR was used to confirm TcII, TcV, and TcVI (300 bp); (3) heminested PCR of the 24S alpha-ribosomal DNA (24Sα-rDNA) was used to distinguish TcV (125 bp) from TcII and TcVI (140 bp); and (4) heminested PCR targeted to genomic fragment A10 was used to discriminate TcII (580 bp) from TcVI (525 bp) [18].

Samples that yielded positive results by SL-IR-II PCR but were non-detectable by 24Sα-rDNA PCR were reported as belonging to the TcII/V/VI group. Those samples that amplified the 140 bp of 24Sα-rDNA fragment but had non-detectable results for A10 PCR were reported as belonging to TcII/VI group. Those samples amplifying both bands of 125 and 140 bp after 24Sα-rDNA PCR, were interpreted as mixed infections by TcV plus TcII and/or TcVI, as previously described [18].

### Statistical analysis

McNemar’s test was used to compare the qualitative qPCR results for S1, S2, and S3 samples from Cochabamba, Tarija, and Aiquile cohorts at baseline, and between baseline and follow-up time-point samples from each treatment group in both clinical trials. The Fisher’s exact test was used to compare the qPCR sensitivity using two or three replicates, and one, two, or three serial samples, and to compare the qPCR positivity between the baseline samples from Cochabamba, Tarija, and Aiquile cohorts, as well as the cumulative therapeutic failure at the end of 12-month follow-up within each treatment group using one, two, or three serial samples, and between BZN arms from both trials. Kruskal-Wallis non-parametric analysis of variance was used to compare the medians of the parasitic loads of quantifiable samples from Cochabamba, Tarija, and Aiquile cohorts at baseline, and from each treatment group at baseline and follow-up time-points. The Tukey’s criterion was used to detect samples with outlier Ct values of IAC (Cts> 75th percentile + 1.5 x interquartile distance of median Ct) [19]. All analyses were performed using SPSS Statistics for Windows V17.0 (SPSS, Chicago, IL).

## Results

### Screening of pre-treated chronic CD patients in DNDi-CH-E1224-001 and MSF-DNDi PCR sampling optimization studies

#### Analysis of qPCR replicates in the DNDi-CH-E1224-001 trial

In this trial, qPCR was firstly assayed in duplicate from each S1 and S2 DNA extract. When both replicates gave non-detectable qPCR results from one of these DNA extracts, a third qPCR replicate was analyzed from the corresponding sample. When the third replicate was included, qPCR positivity increased from 54.8 to 60.5% (for S1) and from 53.6 to 59.2% (for S2) in samples collected from the Cochabamba cohort, and from 59.5 to 63.4% (S1) and from 55.3 to 60.7% (S2) in those collected from the Tarija cohort (Table 1, p>0.05).

**Table 1.** Accumulative qPCR findings in pre-treated chronic Chagas disease patients from E1224 and MSF-DNDi PCR sampling optimization clinical studies, using one, two and three serial samples and two or three qPCR replicates per sample.

| Clinical trial | Locality | Parameters | S1 |  | S2 |  | S1+S2 | S3 |  |
| --- | --- | --- | --- | --- | --- | --- | --- | --- | --- |
|  |  |  | qPCR(1+2) | qPCR(1+2)+3 | qPCR(1+2) | qPCR(1+2)+3 |  | qPCR(1+2+3) | S1+S2+S3 |
| E1224 | CBBA | N | 294 | 294 | 289 | 289 | 294 | 74 | 294 |
|  |  | Positives | 161 (54.8%) | 178 (60.5%) | 155 (53.6%) | 171 (59.2%) | 205 (69.7%) | 19 (25.7%) | 224 (76.2%) |
|  |  | Quantifiables | 31 (19.3%) | 31 (17.4%) | 26 (16.8%) | 26 (15.2%) | 44 (21.5%) | 0 (0.0%) | 44 (19.6%) |
|  |  | Median [IQR] (par. eq./mL) | 2.6 [1.9-3.4] | 2.6 [1.9-3.4] | 2.7 [2.0-3.9] | 2.7 [2.0-3.9] | 2.6 [2.0-3.5] | --- | 2.6 [2.0-3.5] |
|  | Tarija | N | 257 | 257 | 257 | 257 | 257 | 53 | 257 |
|  |  | Positives | 153 (59.5%) | 163 (63.4%) | 142 (55.3%) | 156 (60.7%) | 190 (73.9%) | 10 (18.9%) | 200 (77.8%) |
|  |  | Quantifiables | 37 (24.2%) | 37 (22.7%) | 32 (22.5%) | 33 (21.2%) | 49 (25.8%) | 0 (0.0%) | 49 (24.5%) |
|  |  | Median [IQR] (par. eq./mL) | 2.4 [2.0-3.4] | 2.4 [2.0-3.4] | 3.0 [2.4-3.6] | 3.0 [2.2-3.6] | 2.6 [2.0-3.6] | --- | 2.6 [2.0-3.6] |
| Sampling Study | Aiquile | N | --- | 196 | --- | 195 | 201 | 176 | 205 |
|  |  | Positives | --- | 144 (73.5%) | --- | 150 (76.9%) | 171 (85.1%) | 128 (72.7%) | 185 (90.2%) |
|  |  | Quantifiables | --- | 34 (23.6%) | --- | 40 (26.7%) | 51 (29.8%) | 29 (22.7%) | 61 (33.0%) |
|  |  | Median [IQR] (par. eq./mL) | --- | 2.4 [1.9-4.5] | --- | 2.9 [1.9-4.9] | 2.8 [1.9-4.8] | 3.2 [2.0-4.8] | 3.0 [2.0-4.7] |

S1-3: samples 1-3; qPCR1-3: qPCR replicates 1-3; CBBA: Cochabamba; N: number of samples; IQR: interquartile range; par. eq./mL: parasite equivalents in 1 mL of blood.

#### Analysis of serial blood samples

In the DNDi-CH-E1224-001 trial, the comparison of qPCR positivity obtained after testing individual S1 or S2 samples did not give significant differences (Table 1, p> 0.05) but qPCR positivity increased when the cumulative results from S1+S2 were computed; this was observed in both Cochabamba (60.5 vs 69.7%, p< 0.05) and Tarija cohorts (63.4 vs 73.9%, p< 0.05).

When S1 and S2 gave non-detectable qPCR results, a third sample, named S3 was taken seven days later. The analysis of PCR positivity obtained using three serial samples (S1+S2+S3) compared to that obtained from individual samples demonstrated higher sensitivity for both Cochabamba (60.5 vs 76.2%, p< 0.001) and Tarija cohorts (63.4 vs 77.8%, p< 0.001). Finally, qPCR positivity obtained after testing S1+S2 versus that obtained after testing S1+S2+S3 increased by 6.5% (N= 19/294) in Cochabamba and 3.9% (N= 10/257) in Tarija cohorts (Table 1, p> 0.05).

On the other hand, no statistical difference was observed in qPCR positivity by testing individual S1, S2, or S3 samples in the MSF-DNDi PCR sampling optimization study (Table 1, p> 0.05). Computing the cumulative qPCR positivity obtained for S1+S2 (85.1%) in comparison to the positivity obtained for S1 (10 mL of blood, 73.5%, p< 0.01) or S2 alone (5 mL of blood, 76.9%, p< 0.05) increased sensitivity. This was also true of the cumulative qPCR positivity obtained for S1+S2+S3 (90.2%) compared to that obtained for the individual samples (S1, p< 0.001; S2, p< 0.001; and S3, 72.7%, p< 0.001). Comparison of the cumulative qPCR positivity obtained from S1+S2+S3 with respect to S1+S2 showed an increase of 5.1% (Table 1, p> 0.05).

#### Analysis of T. cruzi DTUs and parasitic loads

It is worth noting the higher qPCR positivity obtained in patients from Aiquile (90.2%) compared to those recruited from Cochabamba (76.2%, p< 0.001) and Tarija (77.8%, p< 0.001); while no difference was found between both E1224 cohorts (Table 1, p> 0.05). Because both studies used the same qPCR method performed in the same laboratory, a hypothesis for this geographical variability in qPCR positivity could be related to diversity of parasitic strains or parasitic loads in the populations studied, and/or to a higher endemicity and exposure to the vector in Aiquile, and therefore a potential risk of reinfection. In order to investigate this, the distribution of *T. cruzi* DTUs was investigated by genotyping 180 qPCR positive samples from these localities with Ct values below 33.

DTUs could be identified in 31 samples: 23 patients were infected with parasite populations belonging to the group TcII/V/VI, six patients were infected with TcI and two presented mixed infections by TcI plus TcII/V/VI (Table 2). TcI was five times more frequent in Cochabamba and Tarija in comparison to Aiquile, although the low number of identified samples preclude determination of its significance. TcIII and TcIV were not detected, not unexpectedly as these lineages have been mostly documented in sylvatic cycle and as a secondary cause of human infection in Venezuela and Brazil [19].

**Table 2.** Direct identification of *T. cruzi* DTUs in blood samples of pre-treated chronic Chagas disease patients from E1224 and MSF-DNDi PCR sampling optimization clinical studies.

| Clinical trial | Locality | *T. cruzi* DTU |  |  |  |
| --- | --- | --- | --- | --- | --- |
|  |  | TcI | TcI+II/V/VI | TcII/V/VI | TcV/VI |
| **E1224** | **CBBA-Tarija** | 5 | 1 | 10 | 1 |
| **Sampling Study** | **Aiquile** | 1 | 1 | 11 | 1 |

DTU: Discrete Typing Unit; CBBA: Cochabamba.

The parasitic loads of baseline samples from the three different cohorts are shown in Fig 1. In Aiquile, 33.0% of samples had parasitic loads above the qPCR LOQ of 1.53 par. eq./mL, whereas in Cochabamba and Tarija the percentage of quantifiable samples was 19.6% and 24.5%, respectively (Table 1). The median and interquartile range values of the quantifiable parasitic loads were 2.6 [2.0-3.5], 2.6 [2.0-3.6], and 3.0 [2.0-4.7] par. eq./mL, for Cochabamba, Tarija, and Aiquile cohorts, respectively (Table 1, p> 0.05).

**Fig 1.**
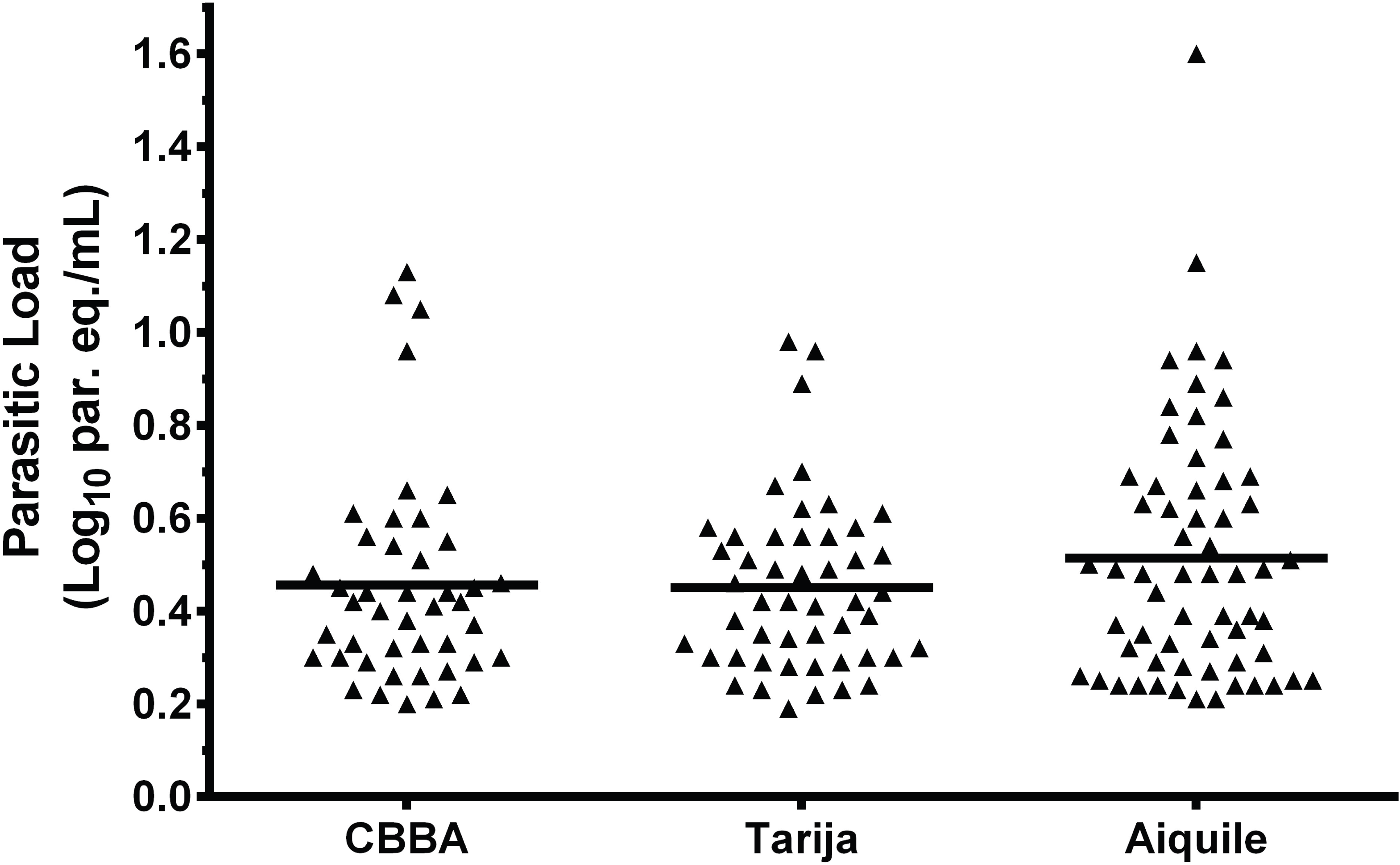
Distribution of parasitic loads in peripheral blood samples of pre-treated chronic Chagas disease patients from E1224 and Sampling Study clinical trials. par. eq./mL: parasite equivalents in 1 mL of blood; CBBA: Cochabamba.

### Follow-up of treated chronic CD patients in DNDi-CH-E1224-001 and MSF-DNDi PCR sampling optimization studies

#### Analysis of qPCR positivity and parasitic loads

Table 3 shows the cumulative qPCR findings obtained from all three serial blood samples during screening and monitoring of all treatment branches in both clinical trials.

**Table 3.** qPCR findings during baseline and follow-up of the different groups of treatment of E1224 and MSF-DNDi PCR sampling optimization clinical studies.

| Clinical trial | Group of treatment | Parameters | BL<br>(S1+2+3) | 2M<br>(S1+2+3) | 4M<br>(S1+2+3) | 6M<br>(S1+2+3) | 12M<br>(S1+2+3) |
| --- | --- | --- | --- | --- | --- | --- | --- |
| E1224 | Placebo | N | 46 | 46 | 46 | 46 | 46 |
|  |  | Positives | 46 (100%) | 34 (73.9%) | 37 (80.4%) | 40 (87.0%) | 36 (78.3%) |
|  |  | Quantifiables | 9 (19.6%) | 15 (44.1%) | 9 (24.3%) | 14 (35.0%) | 16 (44.4%) |
|  |  | Median [IQR]<br>(par. eq./mL) | 2.2 [2.0-4.1] | 2.2 [1.9-4.3] | 3.3 [2.1-4.1] | 3.1 [2.1-3.7] | 2.7 [1.9-5.3] |
|  | E1224 LD | N | 48 | 48 | 48 | 48 | 47 |
|  |  | Positives | 48 (100%) | 5 (10.4%) | 18 (37.5%) | 32 (66.7%) | 36 (76.6%) |
|  |  | Quantifiables | 14 (29.2%) | 0 (0.0%) | 6 (33.3%) | 10 (31.3%) | 12 (33.3%) |
|  |  | Median [IQR]<br>(par. eq./mL) | 3.5 [2.6-7.0] | --- | 2.1 [1.9-2.5] | 2.1 [1.7-2.4] | 2.2 [2.1-4.3] |
|  | E1224 SD | N | 45 | 45 | 44 | 43 | 45 |
|  |  | Positives | 45 (100%) | 4 (8.9%) | 31 (70.5%) | 33 (76.7%) | 38 (84.4%) |
|  |  | Quantifiables | 9 (20.0%) | 0 (0.0%) | 8 (25.8%) | 5 (15.2%) | 12 (31.6%) |
|  |  | Median [IQR]<br>(par. eq./mL) | 2.3 [2.0-2.9] | --- | 2.5 [1.9-3.5] | 2.2 [1.9-2.6] | 2.8 [2.1-4.5] |
|  | E1224 HD | N | 42 | 42 | 41 | 41 | 41 |
|  |  | Positives | 42 (100%) | 7 (16.7%) | 9 (22.0%) | 14 (34.1%) | 23 (56.1%) |
|  |  | Quantifiables | 11 (26.2%) | 0 (0.0%) | 0 (0.0%) | 1 (7.1%) | 6 (26.1%) |
|  |  | Median [IQR]<br>(par. eq./mL) | 2.5 [1.9-3.4] | --- | --- | 1.8 | 2.0 [1.9-2.2] |
|  | BZN | N | 44 | 44 | 43 | 43 | 44 |
|  |  | Positives | 44 (100%) | 3 (6.8%) | 0 (0.0%) | 2 (4.7%) | 2 (4.5%) |
|  |  | Quantifiables | 11 (25.0%) | 0 (0.0%) | 0 (0.0%) | 0 (0.0%) | 0 (0.0%) |
|  |  | Median [IQR]<br>(par. eq./mL) | 2.1 [1.9-2.7] | --- | --- | --- | --- |
| Sampling Study | BZN | N | 137 | 121 |  | 115 | 116 |
|  |  | Positives | 137 (100%) | 28 (23.1%) |  | 11 (9.6%) | 6 (5.2%) |
|  |  | Quantifiables | 47 (34.3%) | 0 (0.0%) | --- | 1 (9.1%) | 0 (0.0%) |
|  |  | Median [IQR]<br>(par. eq./mL) | 2.8 [1.9-4.6] | --- |  | 2.2 | --- |

BL: baseline; 2M, 4M, 6M and 12M: 2, 4, 6 and 12 months from the beginning of the study; S1-3: samples 1-3; LD, SD and HD: low, short and high dosages; BZN: benznidazole; N: number of samples; IQR: interquartile range; par. eq./mL: parasite equivalents in 1 mL of blood

The qPCR positivity of the placebo group from the DNDi-CH-E1224-001 clinical trial was significantly higher at baseline (100%, as per study entry criteria) than at the follow-up time-points (2 months, 73.9%, p< 0.01; 4 months, 80.4%, p< 0.01; 6 months, 87.0%, p< 0.05; 12 months, 78.3%, p< 0.01), whereas no differences were found between the follow-up time-points (Table 3, p> 0.05). Out of the patients who received placebo, 27 were persistently qPCR positive, 15 had intermittently positive and non-detectable results, and four became persistently qPCR undetectable during follow-up.

In both trials, the treated cohorts showed a drastic reduction in PCR positivity at EOT. (E1224 LD, 10.4%; E1224 SD, 8.9%; E1224 HD, 16.7%; E1224 BZN, 6.8%; DND*i*-MSF BZN, 23.1%) (Table 3, p< 0,001). In the E1224 treatment arms, qPCR positivity increased during post-treatment follow-up, reaching its highest value at the end of the study (E1224 LD, 76.6%, p< 0.001; E1224 SD, 84.4%, p< 0.001; E1224 HD, 56.1%, p< 0.01), whereas in cohorts treated with BZN, the proportion of qPCR positive cases diminished at the end of follow-up (E1224 BZN, 4.5%, p> 0.05; DND*i*-MSF BZN, 5.2%, p< 0.01).

Interestingly, all treatment arms showed statistically significant differences in the proportion of positive PCRs between baseline and end of follow-up (E1224 LD, p< 0.01; E1224 SD, p< 0.05; E1224 HD, p< 0.001; E1224 BZN, p< 0.001; DND*i*-MSF BZN, p< 0.001).

The number of patients in the placebo group of the DNDi-CH-E1224-001 trial with quantifiable qPCRs results ranged between 14-16 during follow-up; except at 4 months when, as at baseline, nine patients rendered quantifiable qPCR results (Table 3). Out of nine patients enrolled in the placebo group of DNDi-CH-E1224-001 who showed quantifiable parasitic loads at baseline (Table 3), five showed quantifiable parasitic loads throughout the complete period of follow-up, two patients alternated between quantifiable and non-quantifiable qPCR results, and the two remaining showed persistent detectable, but non-quantifiable, qPCR results throughout follow-up.

Figure 2 shows the distribution of parasitic loads in both clinical studies. No significant differences were found in the medians of parasitic loads at baseline and follow-up time-points in refractory patients with quantifiable parasitic loads (Fig 2A, p> 0.05).

**Fig 2.**
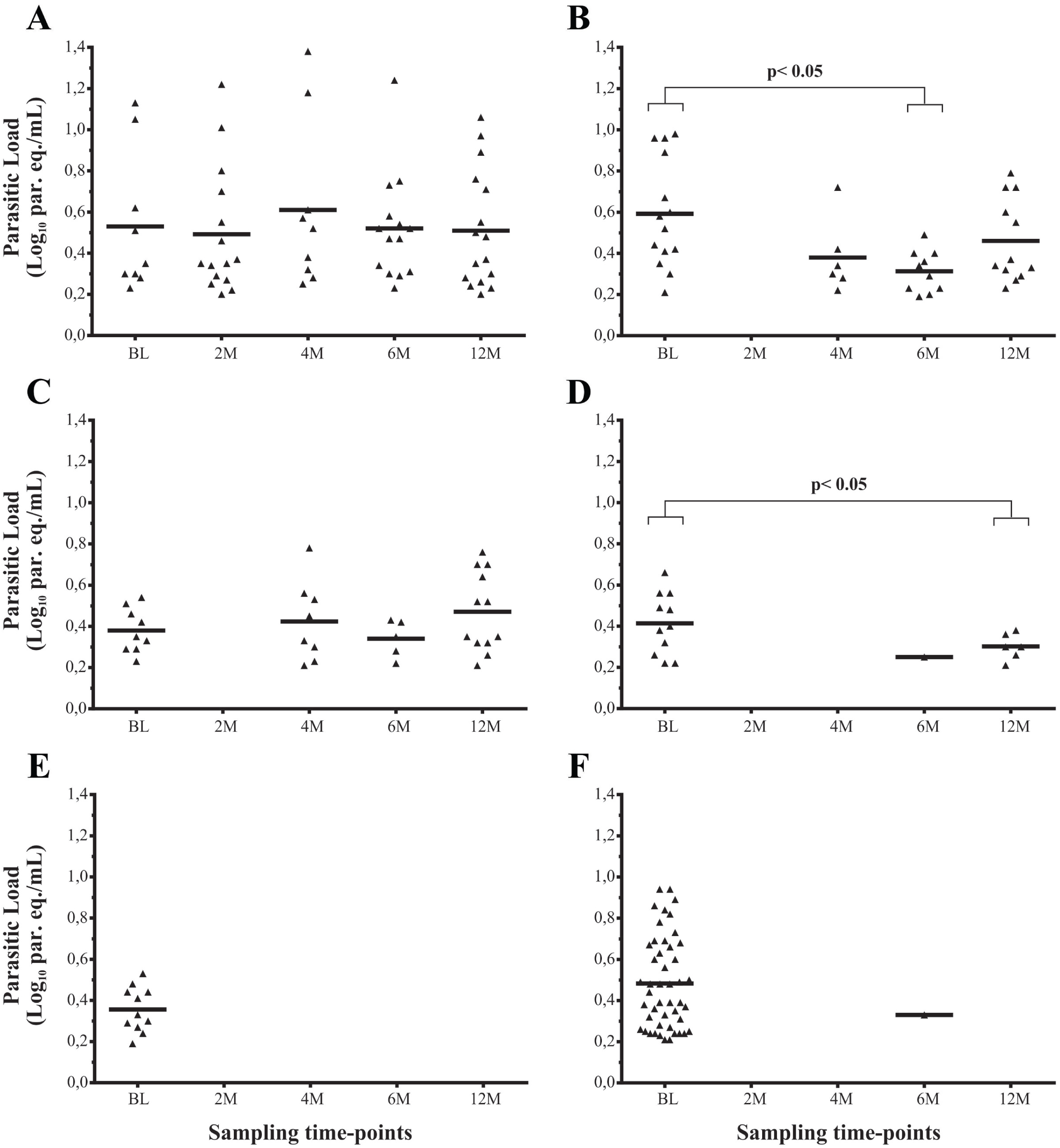
Distribution of parasitic loads during baseline and follow-up of the different groups of treatment of E1224 and MSF-DNDi PCR sampling optimization clinical studies. A: E1224-Placebo arm; B: E1224-Low Dose arm; C: E1224-Short Dose arm; D: E1224-High Dose arm; E: Benznidazole arm from E1224 trial; F: Benznidazole arm from Sampling Study; par. eq./mL: parasite equivalents in 1 mL of blood; BL: baseline; 2M, 4M, 6M and 12M: 2, 4, 6 and 12 months from the beginning of the study

Patients treated with E1224 showed non-quantifiable parasitic loads at the end of treatment, but this increased later on; indeed, 12 cases reached quantifiable loads for E1224 low dose (LD) and short dose (SD) regimes and six in E1224 high dose (HD) regime at the end of follow-up; whereas in BZN treated groups only one sample gave parasitic loads >1.53 par.eq/mL during follow-up (Table 3).

Statistically significant differences were observed between parasitic loads at baseline and 6 months for E1224 LD (3.5 [2.6-7.0] and 2.1 [1.7-2.4] par.eq/mL, respectively; Fig 2B, p< 0.05), and between baseline and 12 months (2.5 [1.9-3.4]) and 2.0 [1.9-2.2] par.eq/mL) for E1224 HD, Fig 2D, p< 0.05).

#### Analysis of cumulative therapeutic failure

Fig 3 compares the cumulative qPCR positivity as a measure of treatment failure obtained for each treatment group in both clinical trials from EOT until the end of follow-up.

**Fig 3.**
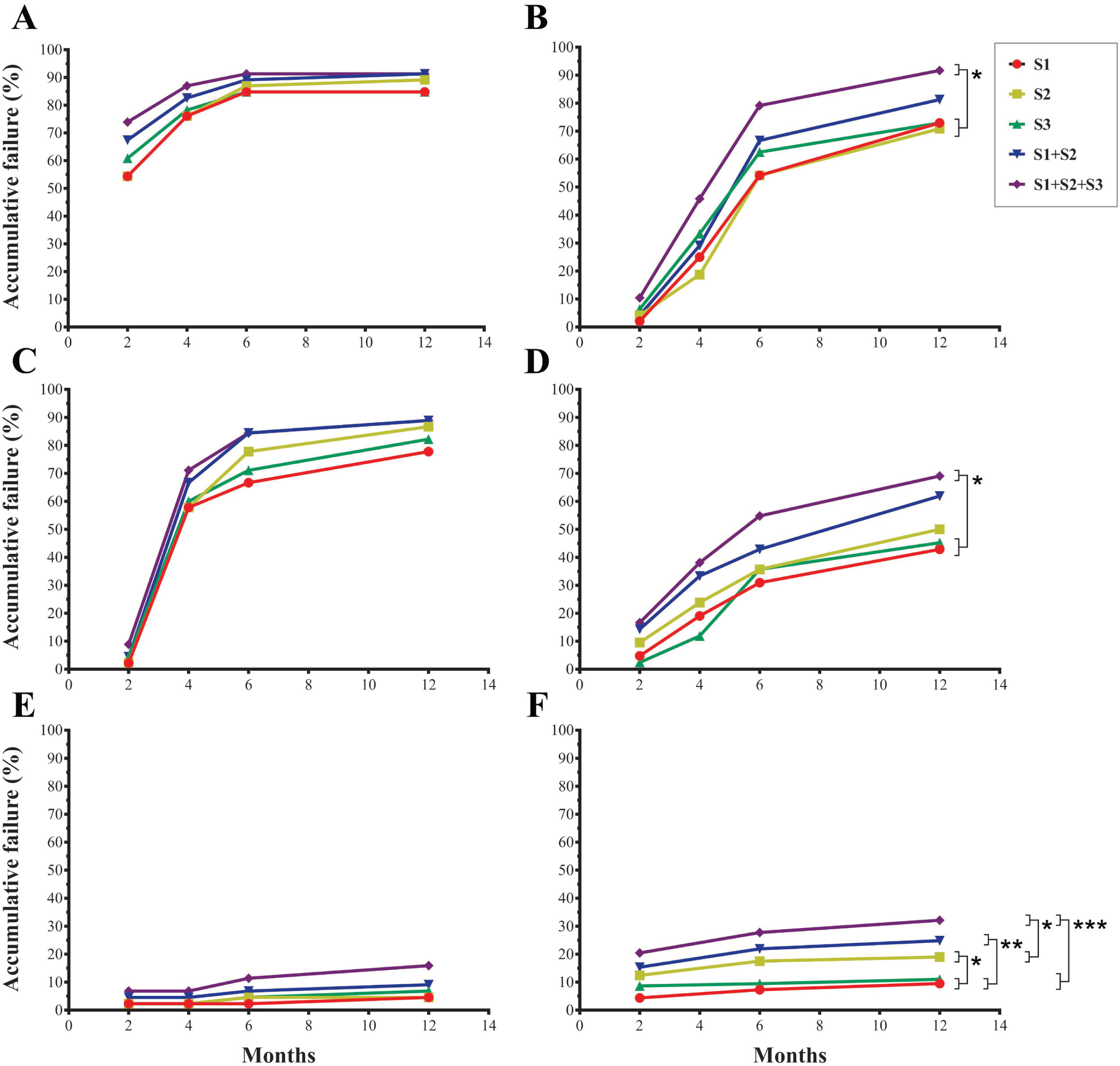
Accumulative therapeutic failure during follow-up of the different groups of treatment of E1224 and MSF-DNDi PCR sampling optimization clinical studies. A: E1224-Placebo arm; B: E1224-Low Dose arm; C: E1224-Short Dose arm; D: E1224-High Dose arm; E: Benznidazole arm from E1224 trial; F: Benznidazole arm from Sampling Study;S1-3: samples 1-3; * p< 0,05; ** p< 0,01; *** p< 0,001

In the DNDi-CH-E1224-001 trial, the multi-sampling strategy (S1+S2+S3) increased detection of treatment failure at the end of follow-up by up to 91.7% for E1224 LD (Fig 3B, p< 0.05), 88.9% for E1224 SD (Fig 3C, p> 0.05), 69.1% for E1224 HD (Fig 3D, p< 0.05), and 15.9% for BZN (Fig 3E, p> 0.05). No significant differences were found between the cumulative treatment failure detected for single S1 (72.9, 77.8, 42.9, and 4.6%), S2 (70.8, 86.7, 50.0, and 4.6%), and S3 (72.9, 82.2, 45.2, and 6.8%) samples, and after testing S1+S2 (81.3, 88.9, 61.9, and 9.1%) versus S1+S2+S3 for E1224 LD, SD, and HD, and BZN, respectively (Fig 3, p> 0.05).

In the MSF-DND*i* PCR sampling optimization study, the strategy involving serial sampling analysis allowed an increase in detection of treatment failure of up to 32.1% (S1+S2+S3) at the end of follow-up in comparison to that detected from individual samples (S1, 9.5%, p< 0.001; S2, 19.0%, p< 0.05; S3, 11.0%, p< 0.001). A significant difference was found in the cumulative treatment failure between S1 and S2 (p< 0.05), whereas no differences were found between S3 and S1 or S2 (Fig 3F, p> 0.05). There was an increase of 7.3% in cumulative treatment failure detected after testing S1+S2+S3 versus that detected after testing S1+S2 (24.8%) (Fig 3F, p> 0.05).

Analysis of cumulative treatment failure tested at EOF in E1224 treatment arms did not show differences between E1224 LD and SD, and between them and the placebo arm (Fig. 3, p> 0.05). In contrast, the E1224 HD arm showed a lower qPCR positivity than LD (p< 0.01), SD (p< 0.05) and placebo (p< 0.05). In addition, the E1224 BZN arm showed lower qPCR positivity than all other arms (p< 0.001).

No statistically significant differences were observed between both BZN treated cohorts enrolled in DNDi-CH-E1224-001 and MSF-DND*i* PCR sampling optimization studies (Figure 3; p> 0.05).

## Discussion

### Impact of serial sampling strategies on qPCR sensitivity

In recent years, several clinical trials to evaluate anti-parasitic treatments for CD were carried out using different sampling strategies and PCR protocols, and variable rates of PCR positivity were obtained [11, 12, 20].

The present analyses shows that qPCR sensitivity was significantly improved at baseline in the DNDi-CH-E1224-001 trial when two blood samples were collected and each DNA extract was analyzed in duplicate by qPCR. The addition of the third blood sample and third qPCR replicate in the subset of patients who had non-detectable PCR results in S1 and S2, gave a small, but non-statistically significant improvement in positivity. The limited data available thus far is insufficient to determine the clinical relevance of this small increase in sensitivity in the evaluation of treatment response. In fact, the samples with only one out of three PCR positive results were non-quantifiable. As treatment was expected to reduce further the parasite burden in those patients with non-quantifiable baseline qPCR results, reducing the chance of detecting treatment failure, three blood samples and qPCR triplicates were tested during post-treatment follow-up.

In the MSF-DND*i* PCR sampling optimization study, the use of 5 mL of blood, instead of 10 ml as starting sample for qPCR analysis, as well as the collection of a third blood sample seven days after the first two samples instead of a few minutes later, did not modify the overall clinical sensitivity [15, 16].

In conclusion, these findings support the use of a lower volume of blood, collected during one visit, for qPCR testing purposes.

### Distribution of discrete typing units and parasitic loads

TcV was the prevailing DTU, in agreement with findings reported by Martinez-Perez et al. 2016 [20], who found TcII/V (64.7 %) to be predominant in Bolivian CD patients living in Madrid, Spain. Differences in qPCR positivity between Cochabamba or Tarija compared with Aiquile could be attributed to a different distribution of parasite DTUs in these localities, such as it was observed with respect to TcI (Table 2), although the low number of identified samples precluded an assessment of the significance of this.

Median parasitic loads were higher in Aiquile than in Cochabamba or Tarija, although the differences did not reach statistical significance (Table 1 and Fig 1). This could be due to the rural nature of the Aiquile area compared to the cities of Cochabamba and Tarija. In a recent study of pregnant women from Bolivia, it was observed that the differences in seroprevalence for *T. cruzi* infection were above all related to the area in which the patients lived most of their lives. Hyper-endemic hotspots were observed where prevalence surpassed 60% and one of the affected areas was the municipality of Aiquile, with 66% seroprevalence [21]. In areas where vector infestation was higher, the seroprevalence of CD was also higher [21].

### Dynamics of bloodstream parasite burden in chronic CD

The monitoring of samples from patients treated with placebo in the DNDi-CH-E1224-001 trial allowed follow-up of the natural history of human chronic *T. cruzi* infection in adult patients for a period of one year. The results showed that a proportion of patients had fluctuations of parasitic loads, which, in some cases, fell below the LOQ (1.53 par. eq./mL) of the qPCR method [10], and even gave non-detectable results, reflecting the fluctuations of parasitaemia observed in chronic patients using traditional parasitological methods [22]. Such findings underscore the need for serial and repeated qPCR examinations for the evaluation of therapeutic failure in chronic Chagas disease.

### qPCR as a surrogate marker of therapeutic failure in CD clinical trials

The qPCR based study of the DNDi-CH-E1224-001 clinical trial demonstrated that BZN was a better parasiticidal drug than E1224 in monotherapy, and that in turn, E1224 HD had higher efficacy than the other E1224 regimens (Fig 2). Treatment with BZN gave a better parasitological response in the urban cohorts of the DNDi-CH-E1224-001 trial than in the rural patients from the MSF-DNDi PCR sampling study, although no significant differences were found. This could be due to the more controlled conditions of treatment administration and follow-up in the DNDi-CH-E1224-001 trial, rather than to a higher risk of re-infection in rural areas, since the houses of patients under treatment were under entomological surveillance.

Finally, this report demonstrates the usefulness of serial blood sampling and performing qPCR replicates not only for enhancing the capacity to recruit chronic CD adult patients to clinical trials, in which the inclusion criteria require a qPCR positive result at baseline, but more importantly for increasing sensitivity to detect treatment failure in this population. It also highlights the importance of standardized methods for monitoring treatment response in chronic Chagas disease.

## Acknowledgements

We thank the patients who took part in this study and the nurses and laboratory staff who contributed to its implementation. We thank Maria de los Angeles Curto (INGEBI-CONICET) for culture of parasite stocks used as positive qPCR controls, constructing standard curves and identification of DTUs. We also thank Dr Louise Burrows (DNDi) for edition of this manuscript. This clinical investigation was funded through DND*i* by the following donors: the Wellcome Trust; Médecins Sans Frontières, International; Ministry of Foreign Affairs, Spain; Department for International Development (DFID), UK; Dutch Ministry of Foreign Affairs (DGIS), Netherlands; Rockefeller Foundation, USA; Federal Ministry of Education and Research (BMBF) through KfW, Germany.. The Platform for a Comprehensive Care of Patients with Chagas disease in Bolivia is a collaborative project between CEADES of Health and Environment; Universidad Mayor de San Simon in Cochabamba, Bolivia; Juan Misael University Saracho, Tarija, Bolivia; and ISGlobal (Barcelona Institute for Global Health, Spain), and supported by the National Chagas Control Program in Bolivia. The Platform is funded by the Spanish Agency for Cooperation and Development (grant number 10-CO1-039). ISGlobal Research group receives funds from the Agència de Gestió d’Ajuts Universitaris i de Recerca (grant number 2014SGR026). JCR is PhD doctoral fellow of CONICET-UBA. AGS is member of CONICET Research Career. FT, JG, IR and AGS are members of NHEPACHA Network.

